# A lung-on-chip model reveals an essential role for alveolar epithelial cells in controlling bacterial growth during early tuberculosis

**DOI:** 10.1101/2020.02.03.931170

**Authors:** Vivek V. Thacker, Neeraj Dhar, Kunal Sharma, Riccardo Barrile, Katia Karalis, John D. McKinney

## Abstract

*Mycobacterium tuberculosis* (Mtb) makes ‘first contact’ with a host in the alveolar space, an interaction largely inaccessible to experimental observation. We establish a lung-on-chip model for early tuberculosis and use time-lapse imaging to reveal the dynamics of host-Mtb interactions at an air-liquid interface with a spatiotemporal resolution unattainable in animal models. By reconstituting host physiology in a modular manner, we probe the role of pulmonary surfactant secreted by alveolar epithelial cells (AECs) in early infection. This is difficult to study directly in animal models, as surfactant-deficient animals are either non-viable or develop acute lung pathologies. We demonstrate that surfactant deficiency results in rapid and uncontrolled Mtb growth in both macrophages and AECs. In contrast, under normal surfactant levels, a significant fraction of intracellular bacteria are non-growing. The surfactant-deficient phenotype is rescued by exogenous addition of surfactant replacement formulations, which have no effect on bacterial viability in the absence of host cells. Surfactant partially removes virulence-associated lipids and proteins ^1,2^ from the bacterial cell surface and consistent with this mechanism of action, we show that attenuation of bacteria lacking the virulence-associated ESX-1 secretion system is independent of surfactant levels. These findings may partly explain why individuals with compromised surfactant function, such as smokers and elderly persons, are at increased risk of developing active tuberculosis.

## Introduction

Early tuberculosis (TB), a respiratory infection caused by *Mycobacterium tuberculosis* (Mtb) is strongly influenced by host physiology; due to the small diameter of respiratory bronchioles only the smallest aerosol droplets containing 1-2 bacilli are successfully transported to the alveolar space ^3^ and the ‘first contact’ with a naïve host is by default a single-cell interaction between an Mtb bacillus and a host cell. There is some evidence that pulmonary surfactant plays host-protective role in these early interactions, but a complete understanding of the role of surfactant is difficult to obtain from animal infection models owing to the lethality of surfactant deficiency. In addition, experiments in animal models ^4^ cannot provide information about the dynamics of host-Mtb interactions at this early stage with sufficient spatiotemporal resolution ^5,6^. A commonly-used *in vitro* model, infection of macrophages with Mtb ^7^, has been used to probe the role of certain surfactant components, but these studies cannot address the role of native surfactant secreted by AECs at an air-liquid interface (ALI), a condition that has been reported to alter Mtb physiology ^8^.

Organ-on-chip systems recreate tissue-level complexity in a modular fashion, allowing the number of cellular components, their identity, and environmental complexity to be tailored to mimic key aspects of the relevant physiology, such as an ALI in a lung-on-chip (LoC) ^9^. These systems have emerged as crucial tools for the replacement of animal models in drug development, toxicity testing, and personalised medicine ^10,11^. A far less-explored line of enquiry has been to use them as models to study the dynamics of host-pathogen interactions in a realistic physiological setting ^12^, where they can combine key advantages of both simpler *in vitro* models and animal models ^13^. Here, we develop a LoC model of early TB infection and use time-lapse microscopy to study the infection dynamics for AECs and macrophages as independent sites of first contact, and the impact of surfactant on infection of AECs and macrophages under ALI conditions that mimic the alveolar environment *in vivo*.

### Lung-on-chip model of early tuberculosis

Freshly isolated mouse AECs comprise a mix of type I cells (Fig. 1A) and type II cells that produce normal surfactant (NS) levels (Fig. 1B). Prolonged *in vitro* passage causes AECs to adopt a phenotype with deficient surfactant (DS) levels (Fig. 1C), reduced expression of type I and type II markers (Fig. 1D), and fewer and smaller lamellar bodies (Fig. 1E, F). We reconstituted a LoC with confluent monolayers of NS or DS AECs (Fig. 1H) and endothelial cells (Fig. 1I) on opposite faces of a porous membrane (Fig. 1J) and an ALI mimicking the alveolar environment. Macrophages added to the epithelial face may remain there or transmigrate across the membrane to the endothelial (Fig. 1H-J). These cells are obtained from a GFP-expressing mouse strain that enables identification during live-cell microscopy. Importantly, NS and DS AECs maintain these phenotypes on-chip at the ALI (Fig. 1K, L), which enables a direct study of the role of AEC-secreted pulmonary surfactant in early infection. Although the ALI significantly degrades axial resolution and signal-to-noise ratios, we are nonetheless able to identify and track individual infected cells over time. Inoculation of the LoC with between 200 and 800 Mtb bacilli led to infection of both macrophages (white boxes in Fig. 1M, P, zooms in Fig. 1O, R) and AECs (yellow boxes in Fig. 1M, P, zooms in Fig. 1N,Q) under both NS (Fig.1M-O) and DS (Fig. 1P-R) conditions. Although the current paradigm in TB focuses on macrophage infection, examination of Mtb-infected cells isolated from the lungs of a mouse at 8 days post infection revealed that 7.3% of infected cells (n = 163) were CD45-pro-SPC+ type II AECs (Fig. S1A-C); we did not find instances of type I AEC infection. The LoC model faithfully reproduces AEC infection, albeit at a higher frequency (Fig. S1D). This in turn enables us to study the infection dynamics in AECs and macrophages simultaneously.

**Fig. 1.**
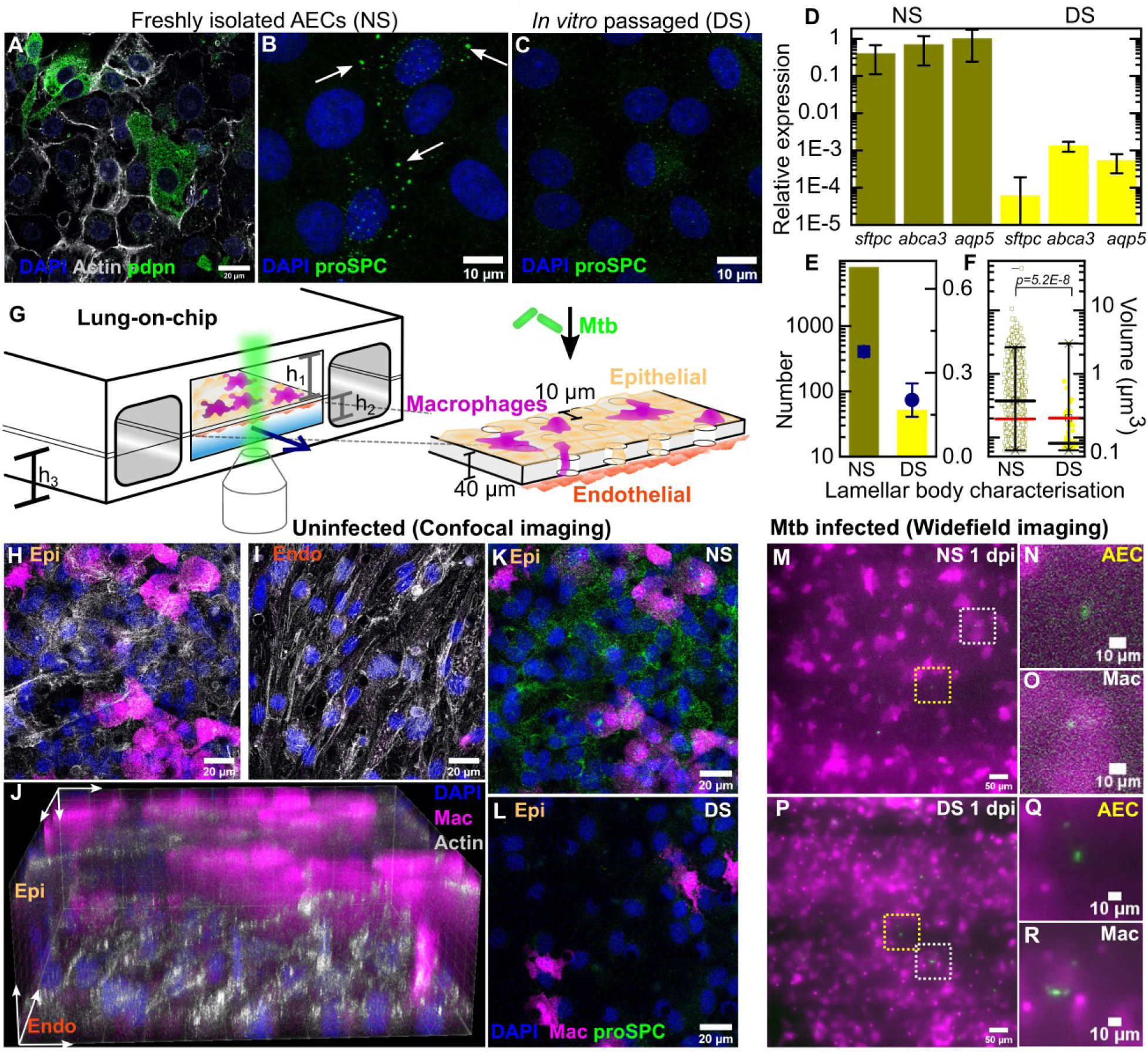
Direct observation of the role of pulmonary surfactant in a lung-on-chip model for tuberculosis. Freshly isolated murine alveolar epithelial cells (AECs) are a mixture of (**A**) type I Pdpn-immunostained cells and (**B**) type II pro-SPC-immunostained cells containing lamellar bodies (white arrows). (**C**) *In vitro* passage of AECs generates an intermediate phenotype with (**D**) reduced expression of type II (*sfptc* and *abca3*) and type I (*aqp5*) markers. Error bars represent the standard deviation for two technical repeats for each qPCR. Based on these characteristics we classify these populations as expressing normal surfactant (NS) or deficient surfactant (DS) levels, respectively. (**E**) In DS cells, the number and mean volume of pro-SPC+ lamellar bodies is reduced. (**F**) The number of lamellar bodies is also significantly reduced. Mean and median volumes are represented by black and red bars, respectively. Whiskers represent the 1-99 percentile interval. (*P* = 5.2E-8) (**G**) Schematic of the LoC model of early tuberculosis. Confluent layers of AECs and endothelial cells populate the top and bottom faces of the porous membrane that separates the air-filled “alveolar” (upper) and liquid-filled “vascular” (lower) compartments, creating an air-liquid interface (ALI). GFP-expressing macrophages (magenta) are added to the alveolar compartment to mimic the natural route of infection. h_1_ = 1 mm, h_2_ = 250 μm, h_3_ = 800 μm. (**H-I**) Confocal microscope images of an uninfected LoC stained to visualize nuclei (blue), actin (grey), and surfactant (green, anti-pro-SPC antibody) verifies that confluency of epithelial (**H**) and endothelial (**I**) layers is maintained at the ALI over 7 days. 3D imaging reveals occupancy of some pores by macrophages (**J**). LoCs reconstituted with NS (**K**) or DS (**L**) AECs continue to express normal or deficient levels of surfactant, respectively. (**M-R**) Widefield microscope images of LoCs reconstituted with NS (**M-O**) or DS (**P-R**) AECs and infected with a low dose of Mtb expressing td-Tomato (green). Images taken at 1 day post-infection (dpi) show that AECs (yellow boxes (**M, P**) and zooms (**N, Q**)) as well as macrophages (white boxes (**M, P**) and zooms **(O, R**)) can be sites of first contact. *P*-values were calculated using a Kruskal-Wallis one-way ANOVA test.

### Surfactant deficiency leads to uncontrolled intracellular growth of Mtb

We used time-lapse microscopy to quantify the intracellular growth of Mtb by measuring the total fluorescence intensity of single infected cells over time (Fig. 2). Between 3 and 5 days post-infection under NS (Fig. 2, A-D, Movie S1) or DS (Fig. 2, E-H, Movie S2) conditions, intracellular growth of Mtb is highly variable in both AECs (Fig. 2I, K) and macrophages (Fig. 2J, L). Plots of the logarithm of total bacterial fluorescence intensity over time for individual infected cells (representing the spread in growth rates) indicate that bacterial growth is exponential in AECs (Fig. 2I, K) and macrophages (Fig. 2J, L). However, under NS conditions, we identified both a substantial “non-growing fraction” (NGF) of bacteria that show very slow growth (doubling time > 168 hours) or even a decrease in fluorescence intensity over time. Intracellular bacterial growth is slower in macrophages compared to AECs under NS conditions but growth rates in both cell types are equivalent under DS conditions (Fig. 2M). Compared to bacterial growth in axenic cultures, growth in both cell types is slower under NS conditions (Fig. S2A) but significantly *faster* under DS conditions (Fig. S2B). Interestingly, there is a much larger spread in growth rates and a small fraction of bacteria continue to grow rapidly even under NS conditions. We observed no spatial pattern of intracellular Mtb growth rates within the lung-on-chip in both NS and DS conditions, confirming that the observed distributions of growth rates are not due to spatial heterogeneity within the device (Fig. S3A, B). Taken together these results suggest that surfactant deficiency shifts the host-pathogen equilibrium in favour of Mtb, resulting in uncontrolled bacterial growth even in macrophages.

**Fig. 2.**
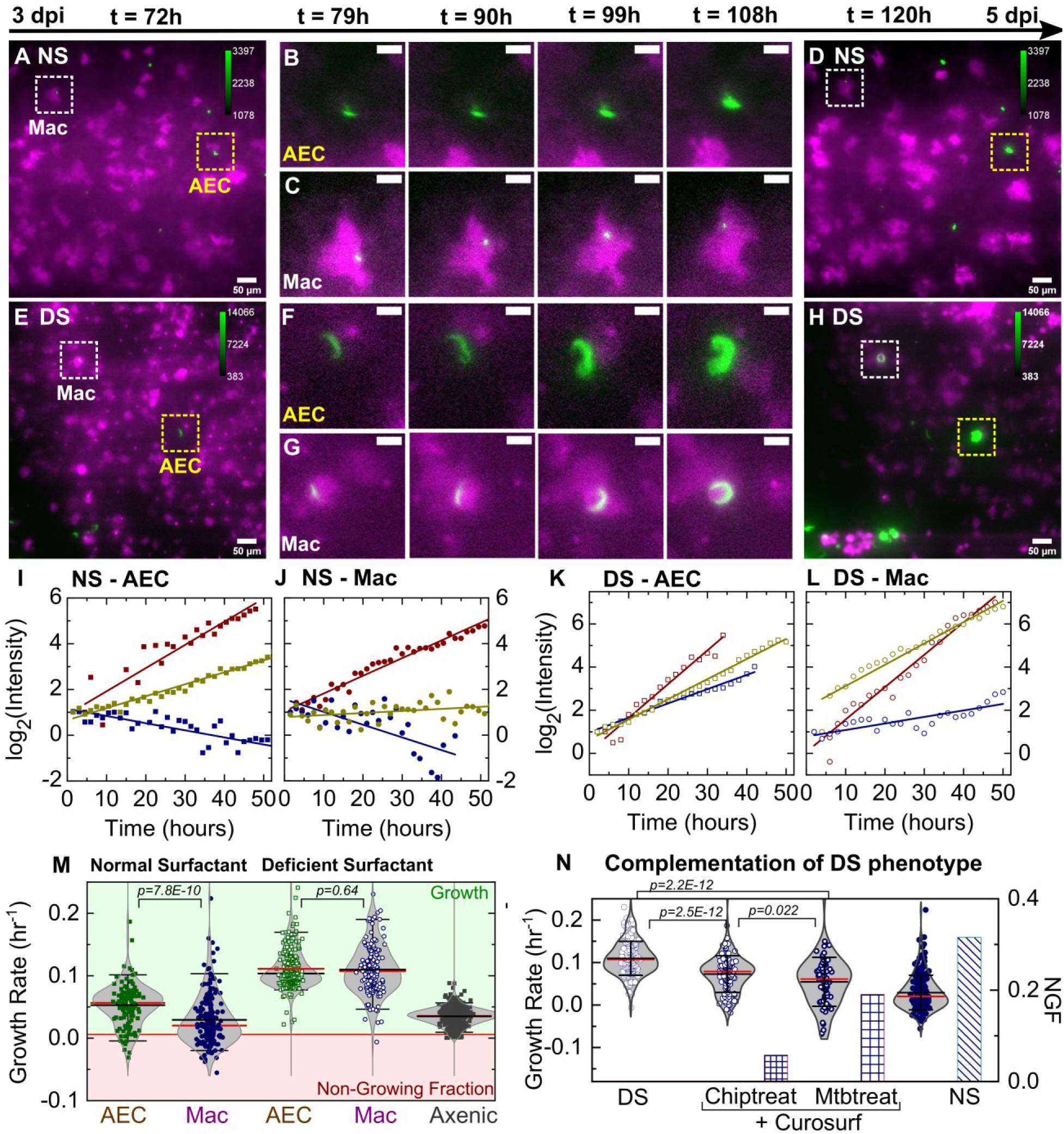
Surfactant deficiency results in uncontrolled intracellular growth of Mtb. Snapshots from live-cell imaging at 1.5- to 2.0-hour intervals between 3-5 days post-infection (dpi) in NS (**A-D**) and DS (**E-H**) LoCs. Macrophages are false-colored magenta; Mtb is false-colored green. The calibration bar (inset in **A, D, E, H**) indicates the absolute intensities in the Mtb channel, the scales were chosen to achieve a similar saturation level in the images across surfactant conditions. Representative examples of infected AEC (yellow boxes) and macrophage (white boxes) are highlighted, and zooms (**B, C, F, G**) reveal growth in both cell types over this period. Scale bar, 10 μm. (**I, J**) Plots of the logarithm of total fluorescence intensity over time confirm exponential Mtb growth for representative infections in AECs (**I, K**) and macrophages (**J, L**) under NS and DS conditions, respectively. In each case, an intracellular microcolony with growth rate close to the population maximum (red), median (yellow), and minimum (blue) is shown. The growth rate is the gradient of the linear fit. (**M**) Scatter plots of Mtb growth rates in AECs (n=122 for NS, n=219 for DS) and macrophages (n=185 for NS, n=122 for DS). Growth is significantly slower in macrophages than AECs in NS conditions (*P* = 7.8E-8) but not in DS conditions (*P* = 0.64), and is more heterogenous in both conditions compared to single-cell Mtb growth rate data from axenic microfluidic cultures. The green- and red-shaded regions indicate growing bacteria and the non-growing fraction (NGF), respectively. (**N**) Uncontrolled growth in DS conditions can be rescued by exogenous administration of Curosurf™. Scatter plots represent Mtb growth rates in a DS LoC treated with Curosurf™ (‘Chiptreat’) or infected with Mtb preincubated with Curosurf™ (‘Mtbtreat’). Data from DS and NS LoC infections (no Curosurf™) are included for comparison. Growth attenuation for both treatments is significant relative to DS conditions as reflected by the average growth rate and the size of the NGF (n=122 for DS, n=121 for Chiptreat, *P* = 2.5E-12 n=63 for Mtbtreat, *P* = 2.2E-12, and n=122 for DS). *P*-values were calculated using a Kruskal-Wallis one-way ANOVA test.

### Exogenous addition of surfactant restores control of Mtb growth

Although reduced surfactant secretion in DS LoCs correlates with increased Mtb replication, this shift could reflect other physiological changes that occur during *in vitro* passage of AECs. We therefore asked whether uncontrolled intracellular replication of Mtb in LoCs reconstituted with DS AECs could be rescued by exogenous addition of surfactant. A 1% solution of Curosurf™, a pulmonary surfactant formulation comprising dipalmitoylphospatidylcholine (DPPC) and the hydrophobic surfactant proteins SP-B and SP-C, was used to treat either Mtb or the DS LoC prior to infection. Both procedures attenuated intracellular Mtb growth and generated a non-growing fraction similar in magnitude to infected NS LoCs (Fig. 2N, Table S1). Curosurf™ treatment does not affect Mtb viability *in vitro* in the absence of host cells (Fig. S4), suggesting that surfactant protects by altering the interaction of Mtb with host cells rather than by any direct effect on bacterial physiology. We conclude that uncontrolled intracellular growth of Mtb in DS LoCs is largely attributable to reduced surfactant secretion.

### Attenuation of an ESX-1 deficient strain of Mtb is independent of surfactant

We examined whether Mtb mutants that were previously shown to be attenuated in the mouse model of tuberculosis are also attenuated in the LoC model and whether surfactant plays a role in attenuation. Mtb lacking the isocitrate lyase genes *icl1* and *icl2* grows normally under standard conditions *in vitro* but is incapable of growth in the lungs of mice and is rapidly cleared ^14^. In the LoC model, we found that the Δ*icl1* Δ*icl2* strain is unable to grow in either AECs or macrophages even under the more-permissive DS conditions (Fig. 3C, S5 A-C), indicating that attenuation of this mutant is similar in the LoC and mouse models and independent of surfactant.

**Fig. 3.**
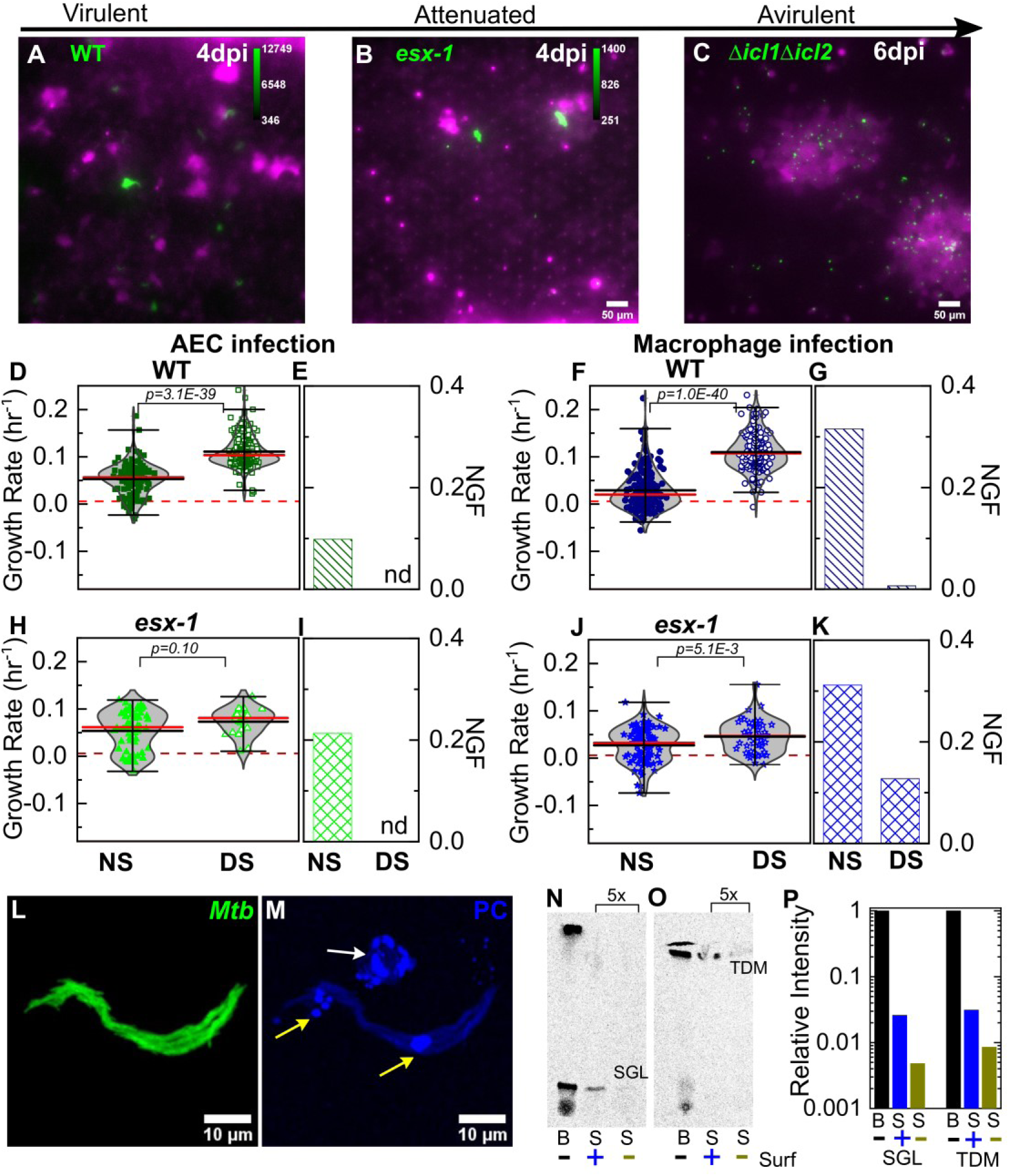
Surfactant depletes virulence factors on the bacterial cell surface. Snapshots from live-cell imaging in DS conditions at 4 days post-infection for (**A**) wild-type Mtb, (**B**) an attenuated ESX-1 deficient strain (*esx-1*), and (**C**) at 6 days post-infection for the avirulent Δ*icl1* Δ*icl2* strain infected at a higher inoculum. The calibration bar (inset in **A-C**) indicates the corresponding absolute intensities in the Mtb channel for a direct comparison across strains. Growth of the ESX-1 deficient strain is attenuated relative to wild-type Mtb (∼10 fold decrease in intensity) and the Δ*icl1*Δ*icl2* strain is unable to grow. Scatter plots indicate Mtb growth rate in individual AECs (**D, H**) and macrophages (**F, J**) and bar graphs indicate the non-growing fraction (NGF) of bacteria for infected AECs **(E, I**) and macrophages (**G, K**) in normal surfactant (NS) and deficient surfactant (DS) conditions for wild-type and ESX-1 deficient strains. ‘nd’, not detected. Each dataset was fitted with a non-parametric kernel density estimation characterised by the mean (black) and median (red) values and whiskers represent the 1-99 percentile interval. For wild-type Mtb, DS conditions significantly increase growth rates and lower the NGF in both AECs (**D, E**) (n=122 for NS, n=219 for DS, *P*=3.1E-39) and macrophages (**F, G**) (n=185 for NS, n=122 for DS, *P*=1.0E-40). For the ESX-1 deficient strain, differences between NS and DS conditions are not significant in AECs (**E**) (n=61 for NS, n=25 for DS, *P*=0.10); for macrophage infection, differences are statistically significant (**J)** (n=93 for NS, n=55 for DS, *P*=5.1E-3), but a significant number of bacteria remain non-growing in DS conditions (**K**). (**L, M**) Maximum intensity projections of an aggregate of fluorescent Mtb (false-colored green, (**L**)) after incubation with a 1% solution of Curosurf™ labelled with 10% v/v TopFluor-Phosphatidylcholine (PC) (false-colored blue, (**M**)). PC is incorporated into surfactant micelles (indicated by arrows in (**M**)), which also associate with and coat the bacteria (yellow arrows). (**N, O**) Thin layer chromatography of total free lipid extracted from wild-type Mtb (B) or bacterial culture supernatants (S) with (+) or without (-) Curosurf™ pre-treatment. The latter two samples were spotted 5x in excess. Running solvents are (**N**) 90:10:1 chloroform: methanol: water to identify sulfoglycolipids (SGL), and **(O)** 80:20:2 chloroform: methanol: ammonium hydroxide to identify trehalose dimycolate (TDM). (**P**) Intensities of the SGL and TDM bands for the three samples in (**N, O**) are plotted relative to that for the bacterial sample without surfactant treatment (labelled ‘B’). *P*-values were calculated using a Kruskal-Wallis one-way ANOVA test.

The activity of the ESX-1 Type VII secretion system, a major Mtb virulence factor that is required for escape from the phagosome into the cytosol ^15^, is upregulated during AEC infection^16^. In comparison to wild-type Mtb, whose intracellular growth rate is strongly dependent on surfactant levels (Fig. 3D-G), intracellular growth of an ESX-1 deficient strain (5’Tn::*pe35* strain ^17^) is largely independent of surfactant levels (Fig. 3H-K) and remains attenuated even under DS conditions. Macrophages are less permissive than AECs for intracellular growth of ESX-1 deficient Mtb under both NS and DS conditions (Fig. S6C, D), in contrast to wild-type Mtb, which grows more slowly in macrophages than in AECs under NS but not DS conditions (Fig. S6A,B). Thus, attenuation of ESX-1 deficient bacteria is not rescued by surfactant deficiency. This demonstrates that ESX-1 secretion is necessary for rapid intracellular growth in the absence of surfactant, consistent with the hypothesis that surfactant may attenuate Mtb growth by depleting ESX-1 components on the bacterial cell surface ^2^.

### Curosurf™ binds to the Mtb cell surface and removes virulence-associated lipids

Although attenuation of the ESX-1 deficient strain is completely independent of surfactant in AECs (Fig. 3H, I), surfactant still has a small but significant impact on growth of ESX-1 deficient Mtb in macrophages (Fig. 3J, K). We therefore hypothesized that an additional mechanism of surfactant-dependent protection could be the removal of virulence-associated lipids from the Mtb cell surface. Consistent with this idea, microscopic examination of Mtb exposed to fluorescently labelled Curosurf™ revealed that surfactant readily coated the bacteria (Fig. 3L, M, S7), albeit heterogeneously (compare Fig, S7A, B with Fig. S7C, D). We also examined the effect of surfactant on the composition of the Mtb cell surface by comparing the total cell-associated lipids and the free (released) lipids prepared from untreated and Curosurf™-treated Mtb. We found that Curosurf™ partially strips the Mtb cell surface of sulfoglycolipids (SGL) (Fig. 3N, P) and trehalose dimycolate (TDM) (Fig. 3O, P), but not phthiocerol dimycocerosates (PDIM) (Fig. S8). The surfactant-mediated removal of these virulence-associated lipids^1^ suggests an additional mechanism for the attenuation of intracellular growth of Mtb in macrophages.

## Discussion

AECs are the major cellular component of the distal lung, yet despite sporadic reports of AEC infection in human tuberculosis ^18,19^, the role of AECs remains controversial. Previous work has revealed the specific role of hydrophilic SP-A and SP-D proteins ^20,21^ and surfactant hydrolase enzymes ^22,23^ on host-Mtb interactions. In contrast, in the LoC model of early tuberculosis presented here, secretion of native surfactant by AECs at an air-liquid interface (ALI) can be modulated. This physiological perturbation, which cannot be achieved in animal models due to the lethality of surfactant deficiency, provides a comprehensive view of the role of AECs in early tuberculosis and is an important advance over previous co-culture models for TB ^24–26^. The unexpectedly rapid and uncontrolled intracellular growth of Mtb at the ALI in the absence of surfactant has not been reported previously in simpler *in vitro* models of host cell infection. Under NS conditions we identified a substantial non-growing fraction of intracellular Mtb in both AECs and macrophages, which may be equivalent to the “non-growing but metabolically active” (NGMA) populations previously observed in the lungs of mice ^27^. These examples underscore some of the advantages of the LoC model, which provides a more faithful mimesis of the complex *in vivo* environment compared to conventional *in vitro* models.

Taken together, our findings indicate that pulmonary surfactant plays on important role in host innate immunity during early Mtb infection, which may partly explain why individuals with defective surfactant function ^28–30^ also show an increased risk of developing active TB. We also report that AECs are more permissive to Mtb growth than macrophages under NS (but not DS) conditions. Infection of AECs is not an artefact of the LoC model, as we also identified infected AECs in the lungs of aerosol-infected mice using a sensitive microscopy-based approach, which might be more discriminating than FACS-based approaches used previously ^31^. Alveolar macrophages in the lung have been shown to be sessile ^5^, and so the higher incidence of AEC infection in the LoC model is probably due to differences in the method of Mtb inoculation. This in turn suggests a deeper link between the airway and alveolar geometries that determine aerosol deposition in the lungs (which are not captured in the LoC model but could be addressed with 3D bioprinted models ^32^) and resident lung immunity. Given the very low inoculum size in human Mtb (most infected individuals harbour just one primary lesion originating from a single bacillus ^33^, we speculate that first contact with an AEC, albeit much rarer *in vivo* could potentially lead to a more aggressive infection. This could provide one explanation for the observation that the proportion of exposed individuals who develop clinical tuberculosis is low.

Time-lapse imaging at an ALI in the LoC infection model directly quantifies bacterial growth rates with a spatiotemporal resolution that is unachievable using indirect measurements of bacterial growth rates in mouse models of tuberculosis (e.g., plating tissue homogenates to measure colony forming units). The deliberate choice to use murine over human cells allows us to add macrophages to the chip from a mouse line that constitutively expresses GFP, which allows unambiguous identification of these cells over days and is superior to the use of CellTrackerTM and other fluorescent dyes. It also enables us to benchmark the model with previous reports from the mouse model for TB. For example, average growth rates for wild-type Mtb under NS conditions are in good agreement with net Mtb growth rates measured using a plasmid-loss assay in mice (Supplementary Discussion) ^34^. However, our microscopy-based approach also reveals the population distribution of growth rates, which highlights that even under NS conditions, a small proportion of cells show robust intracellular Mtb growth, and that robust and attenuated Mtb growth can occur in close proximity to each other on-chip. These observations reinforce an important role for cell-to-cell heterogeneity in host-Mtb outcomes ^35^. The growth characteristics of ESX-1 deficient Mtb in our LoC model are also in good agreement with experiments from animal models, further validating the LoC model (Supplementary Discussion; Fig S9).

Exogenous addition of Curosurf™, a surfactant replacement formulation of phospholipids and hydrophobic proteins, rescues the effects of surfactant deficiency on mycobacterial growth. Phospholipids are the main component of native surfactant, and lipid recycling is a key function of type II AECs and alveolar macrophages ^36^. The two-way interaction of surfactant with the bacterial cell surface (surfactant removes virulence-associated Mtb surface lipids and proteins, and surfactant lipids coat the Mtb surface) may alter how host cells take up these bacteria, or how they are processed after uptake. More generally, surfactant phospholipids have been shown to have antiviral properties, and can serve as a potent adjuvant for antiviral vaccines either on their own ^37^ or as a medium for delivery of immune-stimulating molecules to AECs ^38^. Our findings suggest a potential role for pulmonary surfactant replacement formulations in host-directed therapies against tuberculosis. These insights were made possible by use of an organ-on-chip system that reproduces host physiology in a modular and tuneable fashion, which is frequently impossible to achieve *in vivo*.

## Methods

A detailed experimental protocol for the establishment of the lung-on-chip infection model, a description of the characterization of AECs via qRT-PCR, protocols for live-cell imaging and analyses for quantification of Mtb growth are included in the Materials and Methods in the Supplementary Information. The Materials and Methods also includes an overview of the procedures for the infection of mice with Mtb, generation of single-cell suspension from lungs, and the labelling strategies used to identify infected AECs and immune cells. The Supplementary Information also includes two additional sections of Supplementary Discussion that compare the intracellular growth rate measurements from the lung-on-chip model with those from a plasmid rate-loss assay, and use the growth rate measurements to model the relative attenuation of the ESX-1 mutant versus WT observed using colony forming unit assays.

## Materials and Methods

### Cell culture

Primary C57BL/6 alveolar epithelial cells (AECs) and lung microvascular endothelial cells were obtained from Cell Biologics, USA. Each vial of AECs consisted of a mix of Type I and Type II AECs, which was verified both by immunostaining and qRT-PCR for type I and type II markers (Fig. 1A-C). Both cell types were cultured *in vitro* in complete medium comprising base medium and supplements (Cell Biologics, USA) in 5% CO_2_ at 37°C. Normal surfactant (NS) AECs were seeded directly on the lung-on-chip (see below), without prior *in vitro* culture. Deficient surfactant (DS) AECs were passaged 6-11 times before use.

### Bone marrow isolation and culture

Bone marrow was obtained from 6-8-week-old Tg(act-EGFP) Y01Osb mice (Jackson Laboratories, USA, Stock Number 006567) and cryopreserved. This transgenic line constitutively expresses enhanced GFP under the control of the chicken beta-actin promoter and the cytomegalovirus enhancer. Mice were housed in a specific pathogen-free facility. Animal protocols were reviewed and approved by EPFL’s Chief Veterinarian, by the Service de la Consommation et des Affaires Vétérinaires of the Canton of Vaud, and by the Swiss Office Vétérinaire Fédéral. Bone marrow was cultured in Dulbecco’s Modified Eagle Medium (DMEM) (Gibco) supplemented with 10% fetal bovine serum (FBS, Gibco) and differentiated for 7 days with 20 ng/ml recombinant murine Macrophage-Colony Stimulating Factor protein (M-CSF) (Thermo Fisher Scientific). Bone marrow was cultured in plastic petri dishes without pre-sterilisation (Greiner Bio-one) so that differentiated macrophages could be detached. No antibiotics were used in the cell culture media for all cell types to avoid activation of macrophages or inhibition of Mtb growth.

### Quantitative Real-Time PCR (qRT-PCR)

Freshly isolated AECs (NS) were grown overnight in cell-culture microdishes (Ibidi) or T-25 cell culture flask (TPP, Switzerland). Passaged AECs (DS) were grown to confluency in a T-75 cell culture flask (TPP). Growth media was removed from the flask, and the cells were incubated with the appropriate volume of TRIzol (Ambion) as per the manufacturer’s instruction. TRIzol-treated cell lysates were stored at -20°C before further processing. RNA was precipitated with isopropanol, washed in 75% ethanol, resuspended in 50 μl of DEPC-treated water, treated with Turbo DNase (Ambion), and stored at -80°C until use. DNase-treated RNA was used to generate cDNA using the SuperScript®II First-Strand Synthesis System with random hexamers (Invitrogen), which was stored at -20°C. Specific primers for *gadph, sftpc, abca3*, and *aqp5* are listed in Table S2. qRT-PCR reactions were prepared with SYBR®Green PCR Master Mix (Applied Biosystems) with 1 μM primers, and 2 μl cDNA. Reactions were run as absolute quantification on ABI PRISM®7900HT Sequence Detection System (Applied Biosystems). Amplicon specificity was confirmed by melting-curve analysis.

### AEC characterization via immunofluorescence

Freshly isolated AECs (NS) or passaged AECs (DS) were grown overnight in 35 mm cell-culture microdishes (Ibidi GmbH, Germany). The confluent layer of cells was subsequently fixed with 2% paraformaldehyde (Thermo Fisher Scientific) in phosphate-buffered saline (PBS, Gibco) at room temperature for 30 minutes, washed with PBS, and incubated with a blocking solution of 2% bovine serum albumin (BSA) in PBS for 1 hour at room temperature. The blocking solution was removed and the cells were incubated with the primary antibody (1:100 dilution in 2% BSA solution in PBS) overnight at 4°C. Antibodies used were anti-Podoplanin Monoclonal Antibody (eBio8.1.1 (8.1.1)), Alexa Fluor 488, eBioscience™ (ThermoFisher Scientific), and anti-pro-SPC antibody (ab40879, Abcam). The cell-culture microdishes were washed 3x in PBS, incubated with a fluorescent secondary antibody (Donkey anti rabbit Alexa Fluor 568 (A10042 Thermo Fisher) in a solution of 2% BSA in PBS for 1 hour at room temperature, then thoroughly washed with PBS and incubated with Hoechst 33342 nuclear staining dye (1:1000 dilution, Thermo Fisher Scientific) for 15-20 minutes for nuclear staining. Confocal images were obtained on a Leica SP8 microscope in the inverted optical configuration.

### Bacterial culture

All bacterial strains were derived from *Mycobacterium tuberculosis* strain Erdman and cultured at 37°C. Liquid medium: Middlebrook 7H9 (Difco) supplemented with 0.5% albumin, 0.2% glucose, 0.085% NaCl, 0.5% glycerol, and 0.02% Tyloxapol. Solid medium: Middlebrook 7H11 (Difco) supplemented with 10% OADC enrichment (Becton Dickinson) and 0.5% glycerol. Aliquots were stored in 15% glycerol at -80°C and used once. All strains were transformed with a plasmid integrated at the chromosomal *attB* site to allow constitutive expression of the fluorescent protein tdTomato under the control of the hsp60 promoter. Wild-type (WT) refers to the Erdman strain constitutively expressing tdTomato. The 5’Tn::*pe35* (ESX-1 deficient) strain was generated using transposon mutagenesis ^1^.

### Infection of mice with Mtb

Female C57BL/6 mice (Charles River Laboratories) were housed in a specific pathogen-free facility. Animal protocols were reviewed and approved by EPFL’s Chief Veterinarian, by the Service de la Consommation et des Affaires Vétérinaires of the Canton of Vaud, and by the Swiss Office Vétérinaire Fédéral. Mice were infected by the aerosol route using a custom-built aerosol machine, as described ^2^. Bacteria were grown to exponential phase, corresponding to an optical density at 600 nm (OD_600_) of 0.5, collected by centrifugation at 2850 *g* for 10 minutes, and resuspended in PBS supplemented with 0.05% Tween 80 (PBS-T). The bacterial suspension was subjected to low-speed centrifugation (700 *g*) for 5 minutes to remove bacterial aggregates. The cell suspension was adjusted to OD_600_ 0.1 with PBS-T in a final volume of 20 ml, which was used to infect mice by aerosol. At 1 day post-infection, a group of 4 mice were euthanized by CO_2_ overdose; the lungs were removed aseptically and homogenised in 3 ml of 7H9 medium. Serial dilutions were plated on 7H11 plates containing 100 μg/ml cycloheximide (Sigma) and colonies were counted after 4-5 weeks of incubation at 37°C. The aerosol infection corresponded to a bacterial load of between 60-100 colony-forming units (CFU) per mouse at 1 day post-infection.

### Extraction and quantification of Mtb-infected AECs

At 8 days post-infection, a group of 5 mice were euthanized by an overdose of ketamine/xylazine anaesthetic, and the lungs were washed with PBS delivered via injection through the right ventricle of the heart to remove excess red blood cells. Lungs from each mouse were removed aseptically, minced into small pieces with scissors, and added to 2.5 ml of lung dissociation media reconstituted as per the manufacturer’s instructions (Lung Dissociation Kit – Mouse, Miltenyi Biotec). The lungs were then dissociated using a gentleMACS Octo Dissociator (Miltenyi Biotec). The resulting homogenate was filtered through a 40 μm cell filter, centrifuged for 10 minutes at 300 *g*, and resuspended in alveolar epithelial cell media supplemented with 10% FBS. The homogenate was then plated in 50 mm glass-bottom cell-culture dishes (Ibidi) and incubated for 36-48 hours to allow for epithelial cell attachment. Additional medium was added to each cell-culture dish at 24 hours. The cells were subsequently fixed with paraformaldehyde and stained for immunofluorescence as already described. Antibodies: anti-Podoplanin Monoclonal Antibody (eBio8.1.1 (8.1.1)), Alexa Fluor 488, eBioscience™ (ThermoFisher Scientific) to label Type I AECs, anti-pro-SPC antibody (ab40879, Abcam) followed by secondary antibody staining (Donkey anti rabbit Alexa Fluor 488 (A21206 Thermo Fisher)) to label Type II AECs, and Alexa 647 anti-CD45 antibody (103124, BioLegend).

### Murine lung-on-chip (LoC) model

LoC made of polydimethylsiloxane (PDMS) were obtained from Emulate (Boston, USA). Extracellular matrix (ECM) coating was performed as per the manufacturer’s instructions. Chips were activated using ER-1 solution (Emulate) dissolved in ER-2 solution at 0.5 mg/ml (Emulate) and exposed for 20 minutes under UV light. The chip was then rinsed with coating solution and exposed again to UV light for a further 20 minutes. Chips were then washed thoroughly with PBS before incubating with an ECM solution of 150 μg/ml bovine collagen type I (AteloCell, Japan) and 30 μg/ml fibronectin from human plasma (Sigma-Aldrich) in PBS buffered with 15 mM HEPES solution (Gibco) for 1-2 hours at 37°C. If not used directly, coated chips were stored at 4°C and pre-activated before use by incubation for 30 minutes with the same ECM solution at 37°C. Endothelial cells were cultured overnight at 37°C and 5% CO_2_ in T-75 cell culture flasks, detached with 0.05% Trypsin, concentrated to 5-10 million cells/ml, and seeded on the bottom face of the PDMS membrane. The chip was then incubated for a short period at 37°C to allow the endothelial cells to spread and subsequently seeded with AECs. Freshly isolated AECs were seeded directly from cryopreserved vials received from the supplier. while DS LoCs were seeded from cells cultured overnight at 37°C and 5% CO_2_, in both cases at a concentration of 1-2 million cells/ml. The chip was incubated overnight with complete epithelial and endothelial media in the epithelial and endothelial channels, respectively, under static conditions. The next day, the chip was washed and a reduced medium for the air-liquid interface (ALI) was flowed through the vascular channel using syringe pumps (Aladdin-220, Word Precision Instruments) at 60 μl/hour as described ^3^. The composition of the ALI media used was as described in ^3^ but with an FBS concentration of 5%. The epithelial face was incubated with epithelial base medium with 200 nM dexamethasone (Sigma Aldrich) without FBS supplementation. Flow was maintained over 2-3 days with daily replacement of the medium on the epithelial face (with dexamethasone supplementation). At the end of this period, GFP-expressing macrophages differentiated for 7 days in M-CSF (described above) were detached from the petri dish using 2 mM ethylenediaminetetraacetic acid (EDTA, Sigma Aldrich) in PBS at 4°C, centrifuged at 300 *g* for 5 minutes, and resuspended in a small volume of epithelial cell media without dexamethasone. This solution containing macrophages was introduced onto the epithelial face and incubated for 30 minutes at 37°C and 5% CO_2_ to allow macrophages to attach to the epithelial cells. Medium on the epithelial face was then removed and the chip was maintained overnight at an air-liquid interface (ALI). Chips that successfully maintained the ALI overnight were transferred to the biosafety level 3 (BSL-3) facility for Mtb infection. No antibiotics were used in any of the cell culture media for setting up the LoC model.

### Immunostaining of uninfected LoCs

Uninfected LoCs were maintained at an ALI for up to 7 days after addition of macrophages, during which time ALI medium was flowed through the endothelial channel at 60 μl/hour. After 7 days at the ALI, the chip was fixed for immunostaining as described above; a permeabilization step with a solution containing 2% w/v saponin (Sigma Aldrich) and 0.1% Triton X-100 (Sigma Aldrich) was performed before incubation with the secondary antibody. F-actin on both the epithelial and endothelial face was stained using Sir-Actin dye (Spherochrome) at 1 μM for 30 minutes concurrently with Hoechst staining, as described above.

### Infection of the LoC with Mtb

The chip was assembled into a stage top incubator (Okolab, Italy) prior to infection and flow of medium through the vascular channel was maintained throughout the course of the experiment by use of a syringe pump. A 1 ml aliquot of a culture of Mtb grown to exponential phase (OD_600_ 0.3-0.5) was centrifuged at 5000 *g* for 5 minutes at room temperature, the supernatant was removed, and the cell pellet was resuspended in 200 μl of epithelial cell media without FBS. A single-cell suspension was generated via filtration through a 5 μm syringe filter (Millipore). The single-cell suspension was diluted 100-fold in epithelial media and 30 μl was added to the epithelial channel of the LoC. The infectious dose was measured by plating serial dilutions of the single-cell suspension on 7H11 plates and counting CFU after 3-4 weeks of incubation at 37°C, and varied between 200 and 800 Mtb bacilli. The chip was incubated for 2-3 hours at 37°C and 5% CO_2_ to allow Mtb infection of cells on the epithelial face, after which the solution on the epithelial face was withdrawn. The proportion of bacteria that remained on the chip was estimated by plating serial dilutions of the withdrawn solution on 7H11 plates and counting CFU after 3-4 weeks of incubation at 37°C. The epithelial face was returned to ALI and the inlets of the infected chip were sealed with solid pins as a safety precaution for time-lapse microscopy imaging in the BSL-3 facility.

### Time-lapse microscopy of the Mtb-infected LoC

The LoC was placed in a microscope stage-top incubator and mounted on the stage of a widefield Nikon Ti-2 microscope. The stage-top incubator was connected to a gas mixer (Okolab) to maintain 5% CO_2_ throughout the imaging period. Flow of medium through the vascular channel was maintained throughout this period via the use of a syringe pump. The chip was imaged using a long working distance 20x phase-contrast objective (NA=0.75, Ph2, Nikon) at 1.5-hour or 2-hour imaging intervals. The epithelial face of the chip (where the refractive index differences were highest due to the ALI) was maintained in focus using the Nikon Perfect Focus System. At each timepoint, a Z-stack of 9-10 images with an axial spacing of 10 μm was taken series for a series of fields of view along the length of the chip to account for the dynamic 3D movement of macrophages between both faces, as well as drift in focus over time. Each field of view was ∼660 x 600 μm^2^. Using a Sola SE II light source (Lumencor, USA), macrophages and Mtb were identified through fluorescence emission in the green (macrophages) and red (Mtb) channels using GFPHQ and mcherryHQ 32 mm dichroic filters, respectively. Phase-contrast images were also captured; the poor quality of these images due to the refractive index differences at the ALI serves as a continuous verification that ALI is maintained. All images were captured with an EMCCD camera (iXON Ultra 888, Andor) cooled to -65°C, with an EM gain setting of 300 to allow the sample to be illuminated with a low intensity of incident light in all fluorescent channels with reduced photodamage. Co-localization of the green and red fluorescence signals over a time course was identified as consistent with macrophage infection. Bacteria that did not co-localize with macrophages over time were assumed to infect AECs, which was verified by subsequent immunostaining.

### Data analysis

Images were visualized using ImageJ. Macrophage and AEC infections from each field of view were visually curated by assessing the co-localization of fluorescent signals over time. Smaller stacks of 1-2 microcolonies were assembled. Custom-written software in MATLAB was used to measure the total fluorescence intensity of each intracellular bacterial microcolony which used the nestedSortStruct algorithm for MATLAB (https://www.github.com/hugheylab/nestedSortStruct,GitHub) written by the Hughey lab. Briefly, at each timepoint, the Z-stack with the highest intensity in the fluorescence channel was identified; this image was then segmented to identify the bacterial microcolony; total fluorescence was measured by summing the intensity of all the pixels in this region after subtracting a value for each pixel that represented the average background fluorescence. We chose to measure the total fluorescence intensity because it accounts for both bacterial growth and dilution of the fluorescent protein due to growth (which is slow in a slow-growing bacterium like Mtb). We were unable to measure microcolony volumes accurately using widefield imaging due to poor axial resolution caused by large refractive index differences at the ALI; therefore, we obtained this value from only the Z stack with the highest intensity. Statistical analysis was performed using Origin 9.2 (OriginLabs) and p-values were calculated using a Kruskal-Wallis one-way ANOVA test, with the null hypothesis that the medians of each population were equal.

### Simulations of *in vivo* infections

Growth rate datasets for wild-type and ESX-1 deficient strains of Mtb in NS and DS LoC conditions were fitted with a non-parametric Kernel Smoothed distribution. We simulated a low-dose aerosol infection of 50 bacteria in the alveolar space of n=100 or n=1000 mice, and conservatively assumed that every bacterium interacted with a macrophage upon first contact. Each bacterium was assigned a growth rate picked at random from the Kernel Smoothed distributions and assumed to grow exponentially with these growth rates to generate an intracellular microcolony. The total bacterial numbers in each mouse at 2, 3, 5, 7 and 14 days post-infection were obtained by summing the bacterial counts from each microcolony for each mouse and are shown in Fig. S9E-G. Total bacterial numbers for n=100 mice of WT and ESX-1 deficient strains are shown in Fig. S9.

### Curosurf™ treatment of DS LoCs

Curosurf™ (Chiesi Pharmaceuticals, Italy) was used as a 1% solution in epithelial medium for all LoC experiments. In the case where Curosurf™ was added to a DS LoC, a 1% solution was introduced to the epithelial face after the macrophages were added but before ALI was introduced for 2 minutes, and then removed. The following morning, this procedure was repeated just prior to the addition of the single-cell suspension of Mtb in the manner described above. Alternatively, a 1 ml aliquot of Mtb in exponential phase in 7H9 media was centrifuged at 5000 *g* for 5 minutes, resuspended in 1 ml of cell culture media containing 1% Curosurf^TM,^ and incubated for 10-15 minutes at room temperature. This solution was then centrifuged again at 5000 *g* for 5 minutes and a single-cell suspension of Mtb was generated as described above. Fluorescent labelling of surfactant was achieved by adding TopFluor phosphatidylcholine (10% v/v, Avanti Polar Lipids) to Curosurf™ before dilution in cell culture medium.

### Total free lipid extraction and thin-liquid chromatography (TLC)

Mtb cultures (10 ml each) were grown to stationary phase in 7H9 with 10 μC_i_ of ^14^C-propionate added during exponential phase. Total free lipid extraction from the bacterial pellet, supernatant, and supernatant from bacteria pre-treated with 3% Curosurf™ for 15 minutes at 37°C were extracted as described ^4^. Extracted free lipids were air-dried, resuspended in 2:1 v/v solution of chloroform: methanol, and aliquots were spotted on 5 x 10 cm TLC silica gel 60 F_254_ (Merck). Running solvent was 90:10:1 chloroform: methanol: water for the analysis of sulfoglycolipids (SGL), 80:20:2 chloroform: methanol: ammonium hydroxide for the analysis of trehalose dimycolate (TDM), and 9:1 petroleum ether: diethyl ether for the analysis of pthiocerol dimycocerosates (PDIM) and triacyclglycerols (TAG). The developed TLC plate was exposed to an Amersham Hyperfilm ECl (GE Healthcare) for phosphorescence imaging and visualized with a Typhoon scanner (GE Healthsciences). Intensities of the bands observed were quantified using ImageJ.

## Data availability statement

The datasets generated during and/or analysed during the current study are available from the corresponding authors on reasonable request. Software code used to analyse these datasets will be uploaded to Github prior to publication.

