## Supplementary Text and Figures for "A lung-on-chip model reveals an essential role for alveolar epithelial cells in controlling bacterial growth during early tuberculosis"

**This PDF file includes:**

Supplementary Text
Figs. S1 to S9
Tables S1 to S2
Captions for Movies S1 to S2

**Other Supplementary Materials for this manuscript include the following:**

Movies S1 to S2

**Supplementary Text**

Comparison of Mtb growth rates calculated in this study and using the replication clock plasmid

Table 1 in [5] lists a mean growth rate $r$ = 0.78 and a mean death rate $\delta$=0.41 for Mtb replication during days 1-14 post infection in mice. Thus, the net growth rate equals $r$-δ=0.37, which corresponds to a doubling time (or generation time) of $t_d = \frac{\ln(2)}{r-\delta} \cdot 24$ hours = 45 hours. In the notation of the current manuscript, this converts to a growth rate ($h^{-1}$) of 0.022, which is in good agreement with the mean or median growth rate that we report for macrophage infections with wild-type Mtb in NS conditions.

LoC model in NS conditions accurately reflects *in vivo* Mtb growth dynamics

Although median values of growth rate per hour are similar for wild-type and *ESX-1* strains of Mtb under NS conditions (Fig. S9A, C), subtle differences in the probability density functions for each distribution (reflected in the 1-99 percentile interval in S9 A, C) could nevertheless generate significant differences in population sizes over a few days. This is particularly true for tuberculosis where growth is exponential in early infection, and bacterial numbers are enumerated over weeks or months of infection in the mouse model. To determine if this could account for the attenuation of the ESX-1 strain observed *in vivo*, we simulated the progression of a low-dose mouse infection (infectious dose=50 CFU at 1 day post-infection) using intracellular bacterial growth rates randomly chosen from the growth rate distributions of each strain in macrophages in the LoC model (Fig. 3F, J). At 2 days post-infection, population sizes for the ESX-1 deficient strains (Fig. S9E, n=100, p=2.3 x $10^{-6}$) are already significantly smaller than for wild-type Mtb. The levels of attenuation predicted by this simple model using values from NS (Fig. S9F) but not DS (Fig. S9G) LoC conditions are in good agreement with the experimental data for both mutants from the mouse model in the acute phase of infection [1,6]. These results provide a strong validation that surfactant secretion by freshly isolated AECs in NS conditions in the LoC model provide a better mimic of the native lung environment than DS conditions.

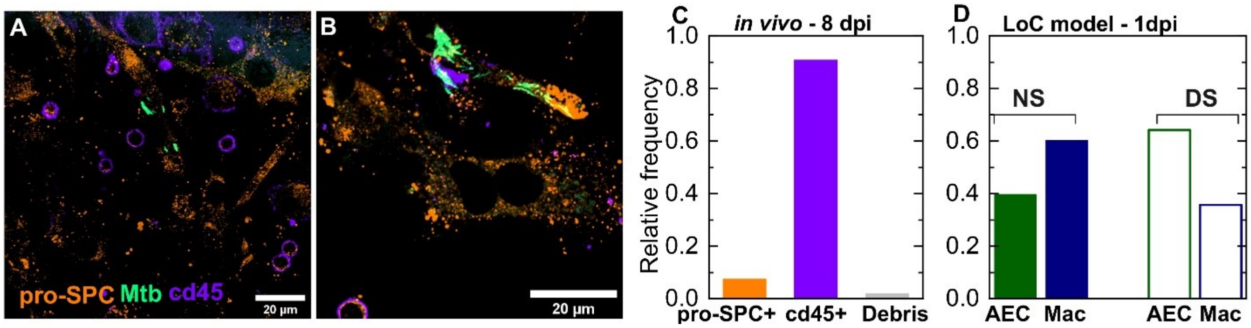


**Fig. S1.**

Examples of a lung cells from a C57BL/6 mouse at 8 days post-infection that show immune cells (indigo, anti-CD45 antibody) and type II AECs (amber, anti-proSP-C antibody) that are (**A**) lightly infected or (**B**) heavily infected. (**C**) Bar plots of the relative frequency of Mtb observed intracellularly in type II AECs, immune cells, or within host cell debris from a total of n=163 infected cells as analyzed by confocal microscopy. (**D**) A bar plot of the relative frequency of AEC and macrophage infections at 1 day post-infection in the LoC model under NS or DS conditions.

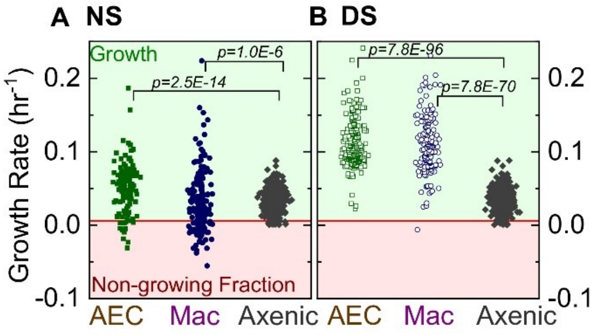


**Fig. S2.**

Scatter plots of bacterial intracellular growth rates for infected AECs (n=122 for NS, n=219 for DS) or macrophages (n=185 for NS, n=122 for DS) in normal surfactant (NS) conditions (**A**) or deficient surfactant (DS) conditions (**B**). These measurements reveal greater cell-to-cell heterogeneity in both conditions (NS and DS) when compared to single-cell Mtb growth rates from axenic cultures grown in microfluidic devices. *P*-values for comparisons of distributions were calculated using a Kruskal-Wallis one-way ANOVA test.

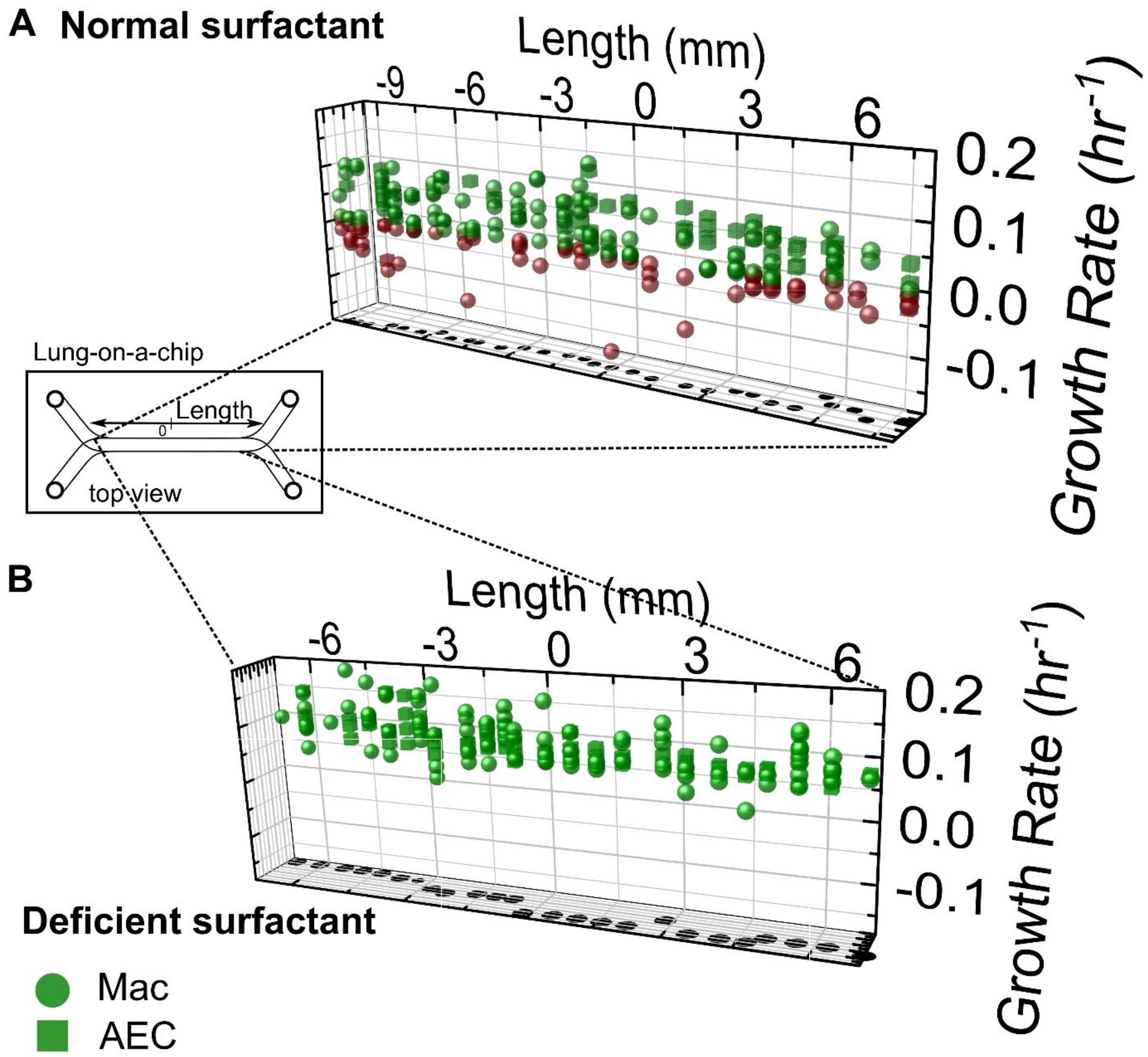


**Fig. S3.**

3D scatter plots of growth rates of individual microcolonies of intracellular bacteria vs. their spatial coordinates on a representative LoC for NS (**A**) and DS (**B**) conditions for infected AECs (cubes) and infected macrophages (spheres). Growing microcolonies are colored green and non-growing fraction (NGF) microcolonies are colored red, as in Fig. 2. In both of the examples shown, heterogenous growth is observed throughout the chip, with no discernable spatial pattern. The X-axis refers to position along the length of the chip, with the approximate position of the origin (0 mm) set roughly in the middle of the chip, as indicated in the schematic.

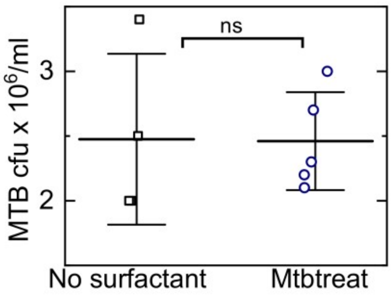


**Fig. S4.**

Exposure to 1% Curosurf™ does not affect Mtb viability in axenic cultures *in vitro*. Single-cell suspensions of Mtb from exponential phase cultures were treated with surfactant (Mtbtreat) or left untreated, and then plated to obtain colony forming units (CFU). The two populations are not significantly different by the Mann-Whitney U Test. Mean values are represented by the black lines, and whiskers represent the standard deviations. Data is obtained from two independent experiments.

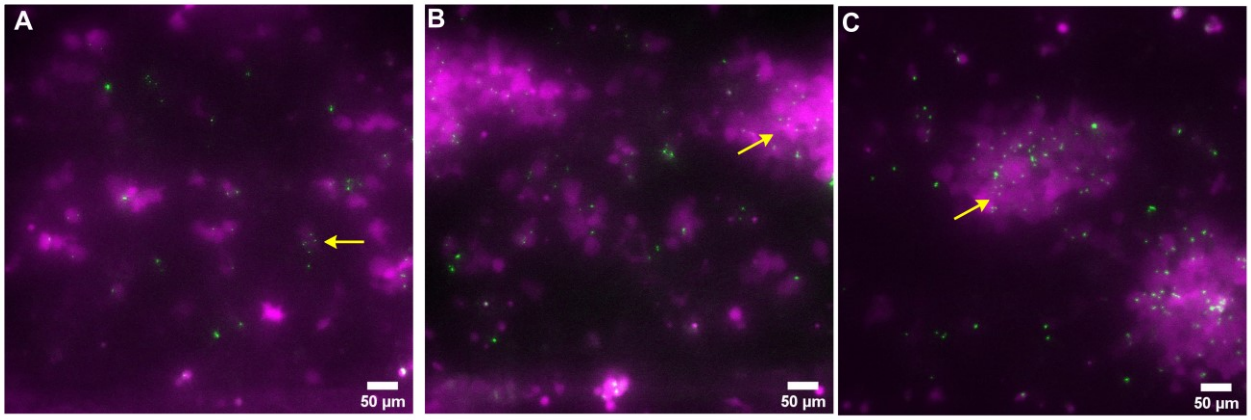


**Fig. S5.**

(**A, B, C**) Snapshots from three representative fields of view taken at 6 days post infection from a DS LoC infected with a high dose of the Δ*icl1* Δ*icl2* double-knockout strain of Mtb. An overwhelming majority of macrophages (false-colored magenta) remain infected with multiple single Mtb bacteria (false-colored green, indicated with yellow arrows) despite the absence of net bacterial growth over time.

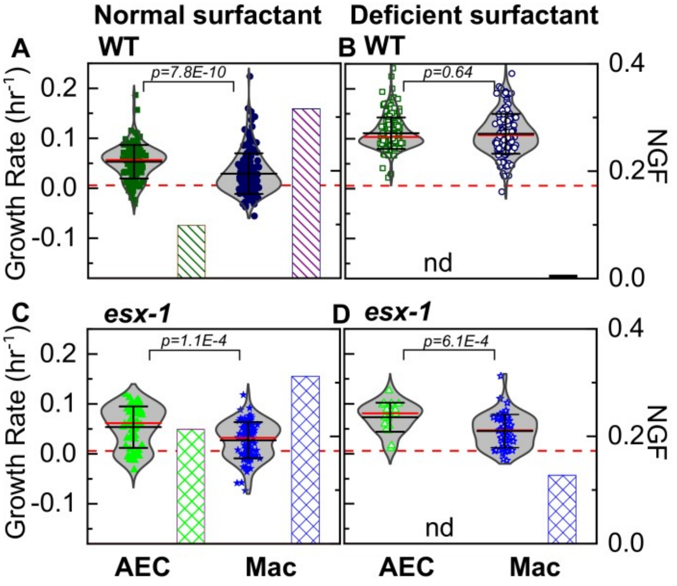


**Fig. S6.**

Scatter plots of growth rates of intracellular bacterial microcolonies and bar charts for the corresponding non-growing fraction (NGF) from Fig. 3 for infected AECs and macrophages under normal surfactant (NS) conditions (**A, C**) and deficient surfactant (DS) conditions (**B, D**) for wild-type and ESX-1 deficient strains of Mtb, respectively. Under NS conditions, AECs are a more permissive niche than macrophages for both strains (**A, C)**. Under DS conditions, macrophages are equally permissive as AECs for Mtb growth for wild-type Mtb (**B**) but not for ESX-1 deficient Mtb (**D**). The number of samples for each strain and condition is given in Table S1. *P*-values were calculated using a Kruskal Wallis one-way ANOVA test.

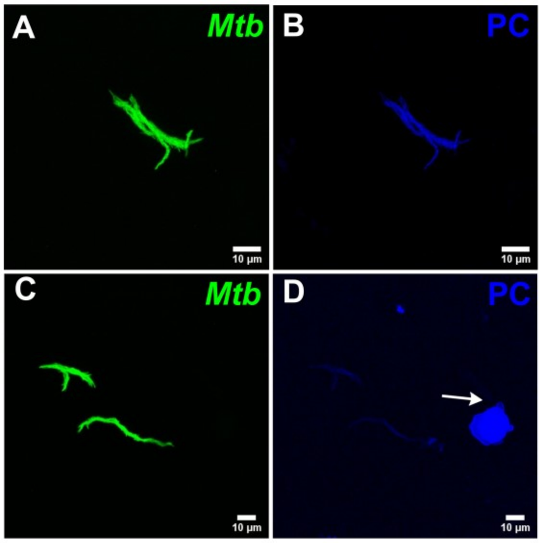


**Fig. S7.**
Additional examples highlighting the interaction of surfactant lipids with the bacterial cell surface. Maximum intensity projections of the bacterial (**A**, **C**) and surfactant (**B**, **D**) channels. A surfactant lipid vesicle is indicated by the white arrow. Intensities in the surfactant channel are normalized across (**B** and **D**) to highlight that surfactant coating of bacteria is heterogenous.

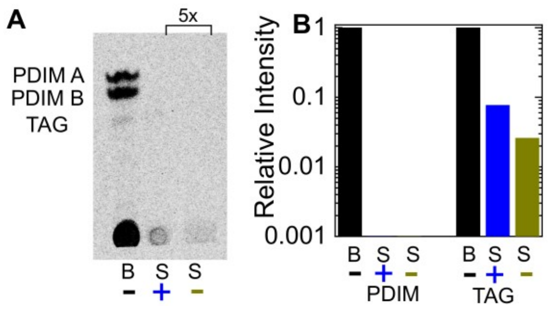


**Fig. S8.**

**A**) Thin layer chromatography of total free lipid extracted from wild-type bacteria (labelled as 'B') or bacterial culture supernatants (labelled as 'S') with (+) or without (-) Curosurf$^{TM}$ pre-treatment. The latter two samples were spotted 5x in excess. Running solvent was 9:1 petroleum ether: dimethyl ether, which identifies phthiocerol dimycocerosates (PDIM) and triacylglycerol (TAG). (**B**) Intensities of the PDIM and TAG bands for the three samples in (**A**) are plotted relative to that for the bacterial sample without surfactant treatment (labelled 'B').

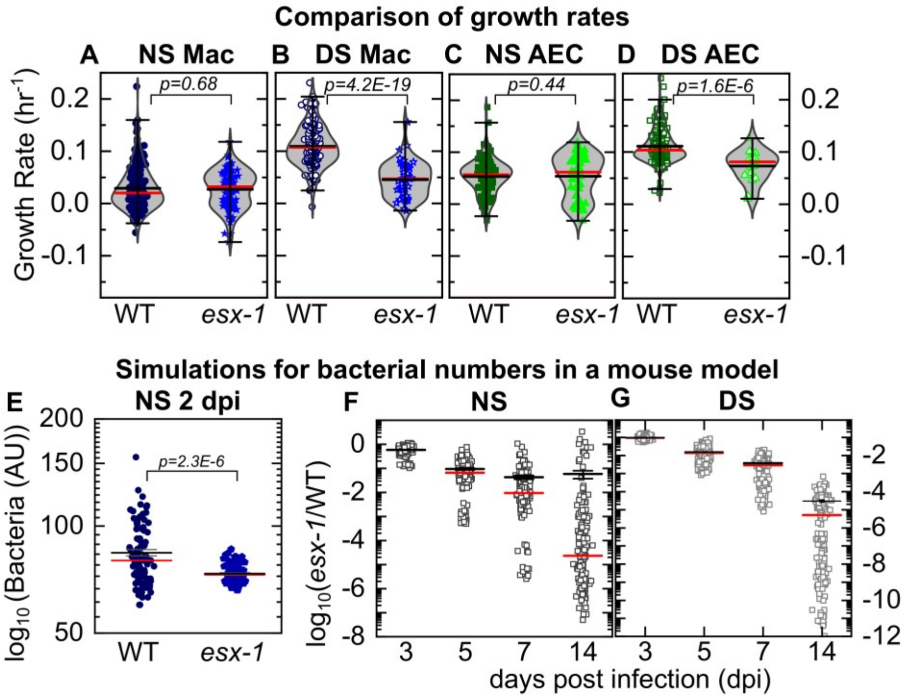


**Fig. S9.**

**Experimental data:** Scatter plots of growth rates of intracellular bacteria from Fig. 3 compared between infected macrophages (**A, B**) and AECs (**C,D**) under NS conditions (**A, C**) and DS conditions (**B, D**) for wild-type and ESX-1 deficient strains (*esx-1*) of Mtb. The number of samples for each bacterial strain and LoC condition is given in Table S1. Mean and median values are indicated by black and red lines, respectively, and whiskers indicate the 1-99 percentile interval. *P*-values were calculated using the Kruskal-Wallis one-way ANOVA test. **Simulations:** Simulations of independent low-dose aerosol infections with 50 WT or *esx-1* bacteria. Simulated bacteria grow at rates randomly chosen from the kernel density estimations for the respective populations in Fig. 3F and 3J respectively. (**E**) In NS conditions, mean bacterial numbers for wild-type Mtb are significantly higher (P=2.3E-6, n=100) than for ESX-1 deficient Mtb. (**F, G**) Plots of the logarithm of ESX-1 deficient population size relative to WT (*esx-1*/WT) at the indicated timepoints for NS (**F**) and DS (**G**) conditions. Each datapoint represents the mean (*ex-1*/WT) ratio from five mice bootstrapped from the larger population (n=1000) for each strain. The attenuation of the ESX-1 deficient strain initially increases but then levels off with a spread of (*esx-1*/WT) ratios by 14 days post-infection. Mean (black) and median (red) values are indicated, and whiskers indicate the standard error of the mean.

| Strain | Infection | Surfactant | Mean($h^{-1}$) | Median ($h^{-1}$) | n |
|---|---|---|---|---|---|
| WT | AEC | NS | 0.053 | 0.056 | 122 |
| WT | Mac | NS | 0.030 | 0.021 | 185 |
| WT | AEC | DS | 0.111 | 0.103 | 219 |
| WT | Mac | DS | 0.109 | 0.107 | 122 |
| *esx-1* | AEC | NS | 0.054 | 0.062 | 61 |
| *esx-1* | Mac | NS | 0.027 | 0.032 | 93 |
| *esx-1* | AEC | DS | 0.073 | 0.081 | 25 |
| *esx-1* | Mac | DS | 0.045 | 0.047 | 55 |
| WT | Mac | DS-chiptreat | 0.073 | 0.079 | 121 |
| WT | Mac | DS-Mtbtreat | 0.055 | 0.061 | 63 |

**Table S1.**

Data for mean and median growth rates and total number of microcolonies (n) analyzed in the different experimental conditions outlined in Fig. 2 and 3.

qPCR primer list
5′-CCGCATCTTCTTGTGCAGTG-3′; *gadph* forward
5′-GATGGGCTTCCCGTTGATGA-3′; g*adph* reverse,
5′-GTAGCAAAGAGGTCCTGATG-3′; s*ftpc* forward
5′-CCTACAATCACCACGACAA-3′; *sftpc* reverse
5′-GCATTGCCCTCATTGGAGAGCCTG-3′; *abca3* forward
5′-TCCGGCCATCCTCAGTGGTGGG-3′; *abca3* reverse

**Table S2.**

Primers used for qPCR characterization of gene expression of the NS and DS AEC cells.

**Movie S1.**

Live-cell imaging over 3-5 days post infection at the ALI for a LoC infected with WT Mtb in NS conditions and corresponding to snapshots in Fig. 2A-D. Imaging frequency is 1.5 hours. Macrophages are false-colored magenta, Mtb is false-colored green.

**Movie S2.**

Live-cell imaging over 3-5 days post infection at the ALI for a LoC infected with WT Mtb in DS conditions and corresponding to snapshots in Fig. 2A-D. Imaging frequency is 2 hours. Macrophages are false-colored magenta, Mtb is false-colored green.

**Biblography**
